# A non-enzymatic acetylation of lysine residues adversely affects the Rubisco activase protein stability

**DOI:** 10.1101/2020.12.21.423885

**Authors:** Li-Li Yang, Hui Hong, Xiang Gao, Jemaa Essemine, Xin Fang, Zhan Shu, Guljannat Ablat, Meng Wu, Hua-Ling Mi, Xiao-Ya Chen, Mingnan Qu, Gen-Yun Chen

## Abstract

The post-translational modifications of non-histone (PTMs) proteins functions are crucial for the plant adaption to the changing environment. The Rubisco activase (RCA) plays a key role in the CO_2_ fixation through the Rubisco activation process. We reported that the RCA from tobacco leaf could be acetylated at several lysine residues including K126 and K164. The acetylation level changes under different light conditions (night and day) as well as under heat stress (45 °C). We further showed that the RCA can be non-enzymatically acetylated *in vitro*, especially by the acetyl-CoA (Ac-CoA) through direct interaction between them. Our results of the *in vitro* assay with deuterium labeled Ac-CoA (D^2^-Ac-CoA) show that the two conserved RCA lysine residues (K126 and K164) were acetylated by Ac-CoA, entraining a dramatic decline in its ATPase activity and a slight effect on the Rubisco activation process. Furthermore, we revealed that the higher RCA acetylation level induced its faster degradation in the chloroplast, which was not a direct consequence of ubiquitination. Eventually, our findings unraveled a new prominent role for the protein acetylation in modulating the RCA stability, which could certainly regulate the carbon assimilation efficiency towards a different energy status of the plants.

## Introduction

Ribulose-1,5-bisphoshate carboxylase/oxygenase (Rubisco), a key photosynthetic CO_2_-fixing enzyme, catalyzes the first reaction of the photosynthetic CO_2_ assimilation, a rate-limiting but notoriously inefficient, step in photosynthesis (Spreitzer and Salvucci, 2002). It forms mostly the inactivated complex with its substrate RuBP and other sugar phosphate inhibitors. The enzyme Rubisco activase (RCA) facilitates the Rubisco activity by the removal of sugar phosphate inhibitors, e.g., RuBP, to maintain a dynamic balance between its deactivation and activation duality (Portis, 2003).

The RCA belongs to the AAA^+^ (an ATPase associated with various cellular activities) family, which is characterized by its prominent ATP hydrolysis activity (Robinson and Portis, 1989). In the presence of Mg^2+^-ATP, a closed-ring hexamers was observed, whereas Mg^2+^-ADP produced amorphous particles (Kuriata *et al.*, 2014). *Nicotiana tabacum* (hereinafter, *N. tabacum*) RCA comprises an N-terminal (N domain) with 67-residues important and crucial for the Rubisco activation process, followed by an AAA^+^ module and a flexible C-terminal extension (Esau et al., 1996). The core AAA^+^ module consists of a nucleotide-binding α/β sub-domain (from helix H_0_ to H_5_) and α helical sub-domain (i-helix H_6_-H_10_) (Stotz *et al.*, 2011; Hasse *et al.*, 2015; Flecken *et al.*, 2020).

The RCA activity responds to different environmental stimuli, such as light and temperature (Chen *et al.*, 2010) and physiological conditions such as source-sink relationship and balance, cellular ionic homeostasis, pH, ADP/ATP ratio (Portis, 2003; Osorio *et al.*, 2014). The RCA thermal stability varies among species, and directly correlates with its photosynthesis thermotolerance threshold (Salvucci and Crafts-Brandner, 2004). Thus, the enhanced RCA thermal stability has been shown to improve the photosynthesis and plant growth under moderate heat stress (Kurek *et al.*, 2007). Moreover, the response of RCA to the oxidoreductive process constitutes an appropriate way to fine-tuning the RCA activity and the Rubisco activation state to the prevailing light intensity. Hence, two Cys-residues in the C-terminal of the large RCA α isoform can reversibly form or break the disulfide bond via thioredoxin-*f in vivo* (Zhang and Portis, 1999). The oxidized RCA, rather than its reduced form, was much more sensitive to the inhibition by the adenosine diphosphate, ADP (Carmo-Silva and Salvucci, 2013). Therefore, the RCA activity was found to be highly sensitive to the ADP/ATP ratios in the stromal chloroplast (Carmo-Silva and Salvucci, 2013).

Some previous reports suggested also that the variation in the RCA content contributes to some extent to the Rubisco activation speed during the changes in the light intensity but not to the overall final activated status of Rubisco (Fukayama *et al.*, 2012; Yamori *et al.*, 2012). The RCA is a thermo-labile protein, as it dissociates, denatures and aggregates under even a moderate heat stress, which could partially be attributed to the disrupted interactions between the RCA monomers and oligomers (Feller *et al.*, 1998; Salvucci *et al.*, 2001). Therefore, the RCA thermal stability varies among species and directly contributes to the photosynthetic efficiency under heat stress (Salvucci and Crafts-Brandner, 2004; Kurek *et al.*, 2007); However, the molecular mechanism underlying the RCA content adjustment and enzymatic regulation under different physiological and environmental conditions remains unclear.

A range of post-translational modifications (PTMs) has been attributed to the plant stress responses (Grabsztunowicz *et al.*, 2017; Zhou *et al.*, 2017; O’Leary *et al.*, 2020). RCA PTMs have been previously well elucidated, such as acetylation (Finkemeier *et al.*, 2011; Wu *et al.*, 2011) and phosphorylation (Kim *et al.*, 2019). Compared to the phosphorylation process, the proteins involved in the photosynthesis typically seems much easier to be modulated by reversible acetylation during the photosynthetic responses to high temperatures (Li *et al.*, 2020). This is due to the fact that Lys-acetylation typically represents a dynamic and reversible PTM occurring mainly on either the α-amino group at the N-terminal or the ε-amino group on the side chain of the Lys-residues (Mischerikow and Heck, 2011).

In this study, we revealed the presence of a dynamic regulation in the RCA acetylation level in tobacco leaf during both day and night times. We further assessed the changes in the RCA acetylation level induced by different metabolites and acetyltransferase together with inhibitors. We investigated as well the crucial relationship of RCA acetylation with its stability, which is required for the higher plants adaption to the changing environmental conditions such as temperature and circadian rhythm.

## Materials and methods

### Plant growth

Tobacco (*N. tabacum*) plants were grown in phytotron under a photoperiod of 12/12h light/dark at 25 (±1°C), light intensity of 450 μmol m^−2^ s^−1^ and a relative humidity (RH) of 65~70%. The Arabidopsis mutant, *sp2* was provided by Prof. Qihua Ling in CAS Center for Excellence in Molecular Plant Sciences. The Arabidopsis were grown in growth chamber at light intensity of 120 μmol m^−2^ s^−1^, temperature 23/22 °C day/night and RH of 70%.

### Total proteins extraction from tobacco leaves

The total soluble proteins were rapidly extracted according to the method as previously described (Chen *et al.*, 2010) with minor modifications. Two histone deacetylase inhibitors (HDACi), 5 mM Nicotinamide (NAM) and 0.2 mM trichostatin A (TSA), were added to the extraction buffer. Firstly, the leaf tissue was grounded using pestle and mortar, then homogenate was transferred into Eppendorf tube, and homogenized in 1:10 protein extraction buffer. The samples were then centrifuged at 14,000 rpm for 10 min at 4 °C. The supernatant obtained via the bicinchoninic acid (BCA) method through chloroplast isolation from tobacco leaves was carried out as described elsewhere (Kubis *et al.*, 2008).

### Extraction and purification of RCA and Rubisco

Tobacco fresh leaves were detached in the morning, then immediately frozen in liquid nitrogen. Thereafter, the RCA was purified from the frozen leaves as reported previously (Chen *et al.*, 2010). The obtained RCA was supplemented with 0.4 mM ATP, and immediately preserved in liquid nitrogen. Rubisco was purified following the protocol as documented previously (Gao *et al.*, 2016). Afterwards, the purified Rubisco was re-suspended in 10% glycerol and preserved at −80 °C, till use. The protein concentration was determined using Bradford protein assay reagent (Bio-Rad, Hercules) followed the manufacturer’s instructions.

### Expression and purification of the recombinant RCA protein

The amino acids substitution for the different RCA sites was carried out on tobacco DNA, and the primers sequences used were listed in Supplementary Table S1. For RCA mRNA and protein sequences, please referred to supporting datasets 1 and 2, respectively. The expression and purification of the recombinant RCA was performed as described earlier (Zhang and Portis, 1999). The site-directed mutagenesis was also achieved through introducing the desired mutation by PCR (Vazyme), followed by the DNA sequencing for confirmation, then cloned into PET28a vector (Novagen). All the constructs were transferred into *E. coli* strain BL21 (DE3) cells, grown at 37 °C for an attenuance at 0.6 optical density (OD) of 600 nm wavelength in Luria-Bertani (LB) medium, followed by induction step with 0.1 mM isopropyl beta-D-thiogalactoside (Sigma) overnight at 20 °C. The cells were then disrupted under French press (Constant systems limited) in a lysis phosphate buffered saline (PBS, pH 7.3), thereafter injected into a fast protein liquid chromatography (FPLC) system (Pharmacia) for purification. Eventually, the purified protein was subsequently stored as described above, till next use.

### Western blot analysis

The standard western-blot procedures were followed in our study; however, for the acetylation approach, the western-blot was performed as reported previously (Wang *et al.*, 2010). The blots were developed with the enhanced chemi-luminescent (ECL) detection system (Thermo Fisher Scientific) and signals were detected and processed using a luminescent image analyzer (LAS-400 mini, GE Healthcare). The levels of RCA protein, lysine acetylation (AcK) and lysine ubiquitin were detected by western blots with anti-RCA (Orizymes), anti-AcK (PTM-105, Jingjie PTM Biolab) and anti-ubiquitin antibodies (MM-0029-P, MEDIMABS), respectively.

### The RCA ATPase activity assay

The RCA ATPase assay was measured by Quanti Chrom TM ATPase/GTase assay kit (BioAsay Systems, DATG-200).

### Acetylation assay of RCA with Ac-CoA, ACP and Ac-AMP

The purified RCA was added to a mixture containing 100 mM Tris-HCl (pH 8.0 at 25 °C), 60 mM NaCl, 0.5 mM Ac-CoA,10 mM ACP or 10 mM Ac-AMP, 1 mM NAM, 1 mM Sodium butyrate, 0.1 mM TSA,2 mM MgCl_2_, 1 mM adenosine triphosphate (ATP). Therefore, 20 μL of the purified RCA (0.5 μg. μl^−1^) was added to 30 μL assay. The reaction mixture was incubated for 30 min at 25 °C before processing.

### Mass spectrum analysis and data processing

The total protein from tobacco leaf was separated in 12% SDS-PAGE and stained with Coomassie brilliant blue (CBB). Thereafter, the protein bands on the gel were sliced, then the gel slices were minced and destained, subsequently the proteins were reduced with dithiothreitol (DTT, Sigma), alkylated with iodo-acetamide (Sigma) and afterwards digested overnight at 37 °C with trypsin (Sigma; in-gel tryptic digestion kit, Thermo Fisher Scientific), desalted with C_18_ Ziptips (Millipore). The resultant peptides were examined by liquid chromatography mass spectrometry (LC-MS). The obtained peptides were identified by LC/MS analysis through Orbitrap and MS/MS.

The MS results were initially investigated through the National Center for the Biotechnology Information (NCBI) database with the aid of the Maxquant software (version 1.4.1.2) as described earlier (Cox *et al.*, 2011). The seeking for the acetylated peptides was carried out against the *N. tabacum* uniprot proteins database (www.uniprot.org). The overall data were processed by Mascot Software (Matrix Science). Accordingly, the peptides were identified by monitoring the maximum false discovery rate (FDR) to be less than 0.01 (< 1%), stricter criteria for the acetylated sites identification which requires a minimum Andromeda peptide score of 60%. An Andromeda is a peptide search engine based on probabilistic scoring. It performs as well as Mascot, can handle data with arbitrarily high fragment mass accuracy, is able to assign and score complex patterns of the post-translational modifications.

The mass recalibration was performed using high confidence identifications based on an initial “first search” using a 20 parts per million (ppm) mass tolerance for parent ion masses and 20 ppm (HCD) or 0.5 Dalton (CID) for fragment ions. Spectra were subsequently sought with a mass tolerance of 6 ppm for parent ions and 20 ppm (HCD) or 0.5 Dalton (CID) for fragment ions. Thus, with the Thermo-Fisher Fusion Orbitrap mass spectrometer, we can perform both CID (collision-induced dissociation) and HCD (higher energy C trap dissociation) collision-induced dissociation studies. In our current investigation, we considered and presented uniquely the acetylated sites and/or peptide with a score more than 40%.

### Surface plasmon resonance analysis

The kinetic studies of the interaction between the Rubisco activase and the substrate Ac-CoA were performed by the surface plasmon resonance (SPR) method using a Biacore T100 instrument (GE Healthcare) at 25 °C as described (Graciet *et al.*, 2004). Thus, the purified RCA protein was immobilized on a carboxy-methylated dextran sensor chip (CM5) chip (GE Healthcare) using an amine coupling method kit (GE Healthcare). A blank immobilization was performed in following the same method and was taken as the reference surface. The RCA was dialyzed against PBS buffer and used as a coupling analyte to the surface of CM5 in the binding assay, then injected over the flow cells at a flow rate of 30 μl min^−1^ for 60 s. The association of the Ac-CoA was monitored for a time period of 120 s, followed by waiting for 180 s, subsequently the disassociation was monitored by the PBS buffer for 420 s. Finally, the regeneration of the surface was performed with a short pulse of 50 mM NaOH. The interaction between RCA and the Ac-CoA was detected and displayed as a sensogram by plotting resonance (RU) units against time. All datasets were analyzed with Biacore evaluation software.

### The RCA degradation assay in vitro

The intact tobacco chloroplast was equally sampled in 24-well plate in 1.5 ml PBS buffer, treated with different inhibitors or acetyl donors for 0, 3, 6 h, then the chloroplast was harvested and detected by western-blot analysis with anti-RCA, anti-AcK and anti-ubiquitin antibodies raised against RCA, AcK and ubiquitin, respectively.

### The Rubisco activation assays

The RCA acetylation by 0.5 mM Ac-CoA was performed as described in detail above. The RCA activity was also assayed as reported earlier (Portis, 2003). Briefly, the purified Rubisco from tobacco leaves was inactivated by desalting on a HiTrap Desalting columns in CO_2_-free buffer containing 20 mM Tricine pH 8.0, 150 mM NaCl and 0.2 mM EDTA, followed by the addition of 1 mM RuBP to form the Rubisco-RuBP complex. The activation process was triggered in a buffer containing 100 mM Tricine pH 8.0, 5 mM NaH^14^CO_3_, 5 mM MgCl_2_, 50 mM DTT, 1 mM ATP, 3 mM creatine phosphate, 14 U ml^−1^ creatine phosphokinase. Therefore, the Rubisco activation was quantified by comparing the initial rate of activation during the first 6 min of the reaction.

### Statistical analysis

A two-tailed Student’s *t*-test was performed for the entire statistical analyses. The data are expressed as the mean of at least three independent experiments (± SD). The differences between samples were assessed at a *P*-value < 0.05(*), *P* < 0.01(**), *P* < 0.001 (***) which mean significant, very significant and strongly significant, respectively.

## Results and discussion

### Temperature and diurnal changes affect acetylation levels of RCA

To compare the RCA acetylation level in tobacco leaves between dark and light conditions, we performed the western-blot analysis with anti-acetylated lysine residue (AcK). The protein acetylation level at ~42 KD, the same molecular weight as RCA, appears oscillating between the day and night times. Thus, the acetylation level was the lowest at noon compared to the other daytime points (Fig. 1A). Meanwhile, the RCA protein amount evaluated by western-blot was also oscillating during the day (Fig. 1A). Notably, the amount of total RCA was the lowest early morning at dawn before the light turning on, and the highest at noon. Furthermore, the acetylation registered at the molecular weight of RCA (42 KD) increased in heat-treated tobacco leaves for 30 min to 45 °C (Fig. 1B), while the RCA amount decreased obviously. In addition, the RCA acetylation level was gradually increased following the increase in the RCA protein amount, especially in stroma-based assays. This suggests that the RCA acetylation level is a temperature- and light-dependent process.

**Fig. 1.**
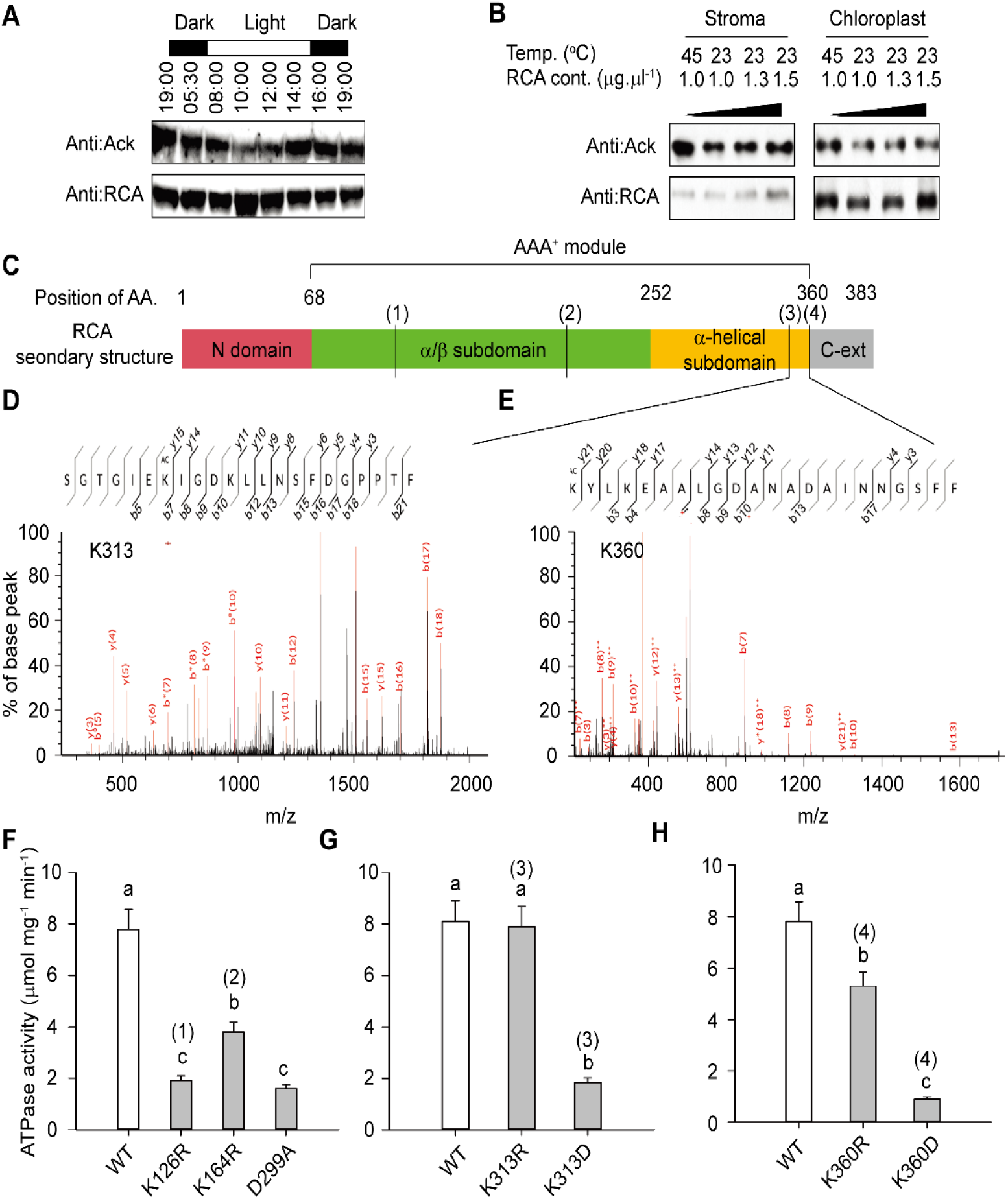
Dark and high temperature increase Rubisco activase (RCA) acetylation in tobacco. (**A**) RCA acetylation level changes in a circadian pattern. The tobacco plants were grown under a photoperiod of 11/13h light/dark. Three-week old fully expanded tobacco leaves were collected in the dark period and 2 h interval during the daytime. For each sample, three leaves were pooled together. In total, around ~3 μg of soluble proteins were loaded for each sample. For the western-blot analysis, the antibodies raised against acetylation and the RCA were used in dilution of 1:10,000. The acetylated tobacco RCA was examined by western-blot with an anti-AcK. The RCA levels were estimated as the signal strength (band size) obtained with the RCA antibody. (**B**) The RCA acetylation level increases at high temperature (45 °C). Fully expanded tobacco leaves were treated for 30 min at either 23 °C (control, CK) or 45 °C, therefore, intact chloroplasts were isolated, and subsequently the total chloroplastic proteins were extracted by SDS protein extract buffer. The total soluble proteins in stroma were also isolated from the intact chloroplast. Then, the acetylated tobacco RCA was examined by western-blot with an anti-AcK. The RCA levels were estimated as well by western-blot using the RCA specific antibody. (**C**) A schematic simplified illustration (mapping) of the RCA secondary structure with its different functional domains, including N domain exhibited from 1 to 68^th^ amino acids (AA), α/β subdomain, α-helical subdomain and C-ext domain which were depicted in pink, green, yellow and grey, respectively. The AAA^+^ module includes α/β subdomain, α-helical subdomain (green and yellow regions). **(D-E)** Mass spectrum of the acetylated lysine residues in tobacco RCA. The RCA from tobacco leaves was extracted, followed by separation with SDS-PAGE and digestion. The acetylated lysine residues were thereafter processed (analyzed) by LC-MS/MS and two acetylated residues were identified to K313 (**D**) and K360 (**E**). **(F-H)** RCA site-directed mutations at the acetylation sites changed their ATPase activities. K126R, K164R and D299A **(F)**, K313R and K313D **(G)**, K360R and K360D **(H)** RCA proteins were expressed in *E.coli.* The ATPase activity for each mutant was determined and compared to that of the wild type (WT) RCA which was purified following the same protocol as for the mutants. Each bar data represents the average of triplicate assays (±SD). The different alphabetic letters adjacent to the histogram bars reflect the significance level at a *P*-value < 0.05 based on one-way *ANOVA* statistical analysis.

### Acetylated lysine residues of RCA is related to ATPase activity

Afterwards, a subsequent analysis using liquid chromatography-tandem mass spectrometry (LC-MS/MS) has identified four acetylated lysine residues, K126, K164, K313 and K360 in tobacco leaves (Fig. 1C-E; Supplementary Table S2). Among them, K126 and K164 are conserved AA residues, which are located at the nucleotide-binding α/β sub-domain (residue 68-252), whereas K313 and K360 residues are situated on the α-helical subdomain and the flexible C-terminal extension, respectively (Stotz *et al.*, 2011). The K313 residue in *Arabidopsis* RCA has been reported as one of the thermo-stable related sites(Kurek *et al.*, 2007). The functions of the other three lysine residues (K126, K164 and K360) had not yet been reported elsewhere. Our results regarding the ATPase activity assay show that the substitutions of these four lysine residues (K126R, K164R, K313R and K360R) significantly decreased the RCA ATPase activity (*P* < 0.05; Fig. 1F-H).

Besides, the K111 of the Arabidopsis RCA, one of the P-loop region sites of the Walk-A domain, has earlier been reported to be acetylated (Finkemeier *et al.*, 2011). Interestingly, we did not identify acetylation modification of RCA K111 in tobacco leaves, which might be related to low acetylation levels. Consistent with the previous results (Shen *et al.*, 1991), the K111R substitution in the tobacco RCA has also led to a lower ATPase activity (Supplementary Fig. S1). Taken together, the RCA acetylation seems to be tightly related and/or dependent to the hydrolysis of its ATP activity as documented previously (Wu *et al.*, 2011).

### RCA can be acetylated by Ac-CoA in absence of GCN5 and Pat

The acetylation of the metabolic enzymes was obtained to be regulated by the histone acetyltransferase and deacetylase (Wang *et al.*, 2010; Lima *et al.*, 2011). Accordingly, several photosynthesis-related proteins were also found to be acetylated(Jiang *et al.*, 2018; Li *et al.*, 2020), as the RCA is a chloroplastic protein (Portis, 2003); however, no canonical acetyltransferase was reported in *Arabidopsis* chloroplast proteome (Peltier *et al.*, 2006). In this regard, we first attempted to acetylate the RCA *in vivo* using two canonical acetyltransferases reported for animal, the general control non-derepressible 5 (GCN5) and the protein acetyl-transferase (Pat); however, the RCA acetylation level was almost the same after incubation with or without any canonical acetyltransferase (GCN5 or Pat) in the assay (Supplementary Fig. S2A-B). Consistently, the acetylation level of RCA proteins expressed in *Pat* deficient *E. coli mutant* (*pat*) was also nearly similar to that of the wild type (Supplementary Fig. S3). Therefore, we speculated that the RCA acetylation could be orchestrated in a concerted manner by a different biological pathway.

Enormous evidences showed that proteins can be acetylated by small molecular metabolites. In mitochondria, the protein acetylation could be facilitated by the alkaline pH and the high concentrations of Ac-CoA and succinyl-CoA (Wagner and Payne, 2013). Besides, the acetyl-phosphate (ACP) can globally and chemically acetylate the lysine residues of protein in *E. coli* (Weinert *et al.*, 2013). The Ac-AMP could also non-enzymatically acetylate the proteins residues (Ramponi *et al.*, 1975). Our previous study showed that the Rubisco can be directly acetylated by a plant-derived metabolite, 7-acetoxy-4-methylcoumarin (AMC) (Gao *et al.*, 2016).

To investigate the eventual RCA acetylation by these metabolites candidates, we examined the RCA acetylation level following *in vitro* incubation with Ac-CoA, Ac-AMP, AMC and ACP, separately, without acetyltransferases furnished in the assay. Notably, the western-blot analysis shows that all the four metabolites candidates mentioned above could merely acetylate RCA (Fig. 2A-C; Supplementary Fig. S2C). The RCA acetylation level enhances with increasing the concentration of the four metabolites, especially for the Ac-AMP (Fig. 2A-C; Supplementary Fig. S2C). Interestingly, the increased RCA acetylation levels due to the Ac-AMP effects also promote the Rubisco acetylation level (Supplementary Fig. S4A).

**Fig. 2.**
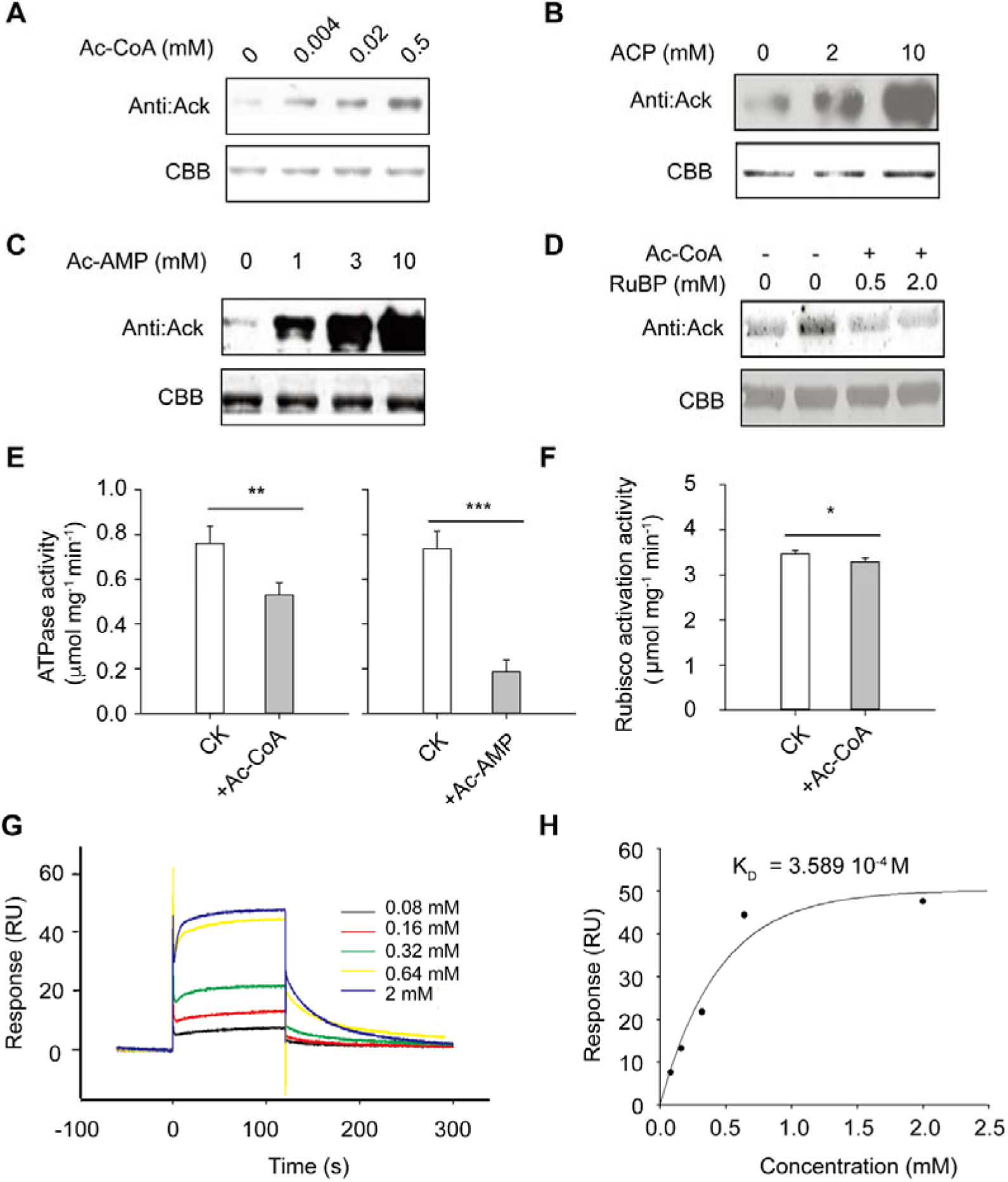
RCA acetylation by different acetyl group donors. **(A-C)** The tobacco RCA protein expressed in *E*. *coli* was acetylated by Ac-CoA (**A**), ACP (acetyl phosphate; **B**) and Ac-AMP (Ac-AMP; **C**). The RCA was firstly incubated with Ac-CoA, ACP and Ac-AMP for 30 min at 25 °C, and then the RCA acetylation level was assessed by western-blot analysis. (**D**) A pre-treatment with RuBP inhibits the RCA acetylation by the Ac-CoA. Indeed, RCA was incubated with RuBP for 5 min at 25 °C before its (RCA) incubation with Ac-CoA (0.5 mM) for 30 min. (**E**) The RCA acetylation by the substrates such as Ac-CoA and Ac-AMP adversely affects the RCA ATPase activity. The RCA was acetylated by 0.5 mM Ac-CoA or 10 mM Ac-AMP, therefore RCA was incubated for 30 min at 25 °C, and then the ATPase activity was determined. (**F**) The Ac-CoA modifies the RCA but did not influence the Rubisco activation process (activity). The RCA was incubated with 0.5 mM Ac-CoA for 30 min at 25 °C and the ATPase initial activity was immediately measured before any potential pre-activation. For panels E and F, each data bar represents the mean of three independent replicates (n = 3) ±SD, and the significance level between the control (CK) and the treated sample was calculated via the Student’s *t*-test, where *P* < 0.01 (**) means very significant and *P* < 0.001 (***) signifies highly significant; ns stands for non-significant (P >0.05). **(G-H)** The interaction between the RCA and Ac-CoA was performed by the surface alkaline resonance method. The RCA was directly immobilized to the sensor chip and the different concentrations of Ac-CoA (0.08 mM, 0.16 mM, 0.32 mM, 0.64 mM and 2 mM) were injected at a flow rate of 20 ul.min^−1^ (**G**). The affinity constant of dissociation (K_D_) was thus determined by simulating the kinetic curves and deduced accordingly (**H**).

It is worthy to mention that the RCA could be acetylated by just 0.004 mM Ac-CoA, much less than its prevailing physiological concentration (0.01-0.02 mM), by up to 5 times in spinach chloroplasts (Post-Beittenmiller *et al.*, 1992). Except the Ac-CoA, the three other metabolites candidates (Ac-AMP, ACP and AMC) are not natural or common metabolites for the higher plants. Therefore, these results suggest that the RCA acetylation seems to be mediated non-enzymatically by Ac-CoA *in vivo*, a major abundant metabolite in the chloroplast.

Furthermore, Ac-CoA and Ac-AMP could dramatically decrease the RCA ATPase activity, but slight decrease the Rubisco activation (Fig. 2E-F; Supplementary Fig. S4B), which remains similar as for the K313 mutation with slight changes in the Rubisco activation process (Kurek *et al.*, 2007). This might be attributable to the fact that the conserved RCA lysine sites in the C-terminal flexible extension and the N-terminal were not modified by the Ac-CoA. These two sub-domains are very important for the Rubisco activation and the truncation of these two sub-domains would lead to a loss in the Rubisco activation properties or peculiarities (Esau *et al.*, 1996).

### The RCA acetylation by Ac-CoA is required for their interaction

As mentioned above, the RCA could be acetylated by the Ac-CoA at a lower concentration than its physiological concentration in the chloroplast. Therefore, the non-enzymatic acetylation method by the Ac-CoA may constitute an important and pioneering biochemical approach for the RCA acetylation *in vivo*. We hence examined the possible interaction between RCA and the Ac-CoA through the SPR. The steady-state affinity data shows that the RU increases when the Ac-CoA concentration increase (Fig. 2G-H), which is consistent with the results of acetylation assay (Fig. 2D). The calculated dissociation equilibrium constant (K_D_) was 0.36 mM; however, when the Ac-CoA was replaced by its analogue malonyl-CoA, the interaction signal was disappeared (Supplementary Fig. S5A-B). The SPR data indicates the existence of a specific interaction between the Ac-CoA and RCA, suggesting the possibility of RCA direct acetylation by Ac-CoA *in vivo*.

### RuBP inhibits the acetylation levels of RCA by Ac-CoA

Previous studies suggested that the content of Ac-CoA was maintained at a stable level during the day time, indicating that the difference in the RCA acetylation level in dark and under light is not ascribable to the changes in the Ac-CoA concentration *in vivo* (Tumaney *et al.*, 2004; Chen *et al.*, 2017). It is actually known that the RuBP has a competitive binding ability with the RCA (Jensen, 2000). In this context, we have previously reported that the Rubisco acetylation could be inhibited by RuBP, a Rubisco substrate with higher expression under light than under dark conditions (Gao *et al.*, 2016). Thus, we expect that this competitiveness binding capacity between them (RuBP and RCA) could inhibit the acetylation process by the Ac-CoA, thus leading to a decrease in the RCA acetylation level. We then examined the RuBP effect on the RCA acetylation process by the Ac-CoA, and confirmed its inhibitory effects on RCA acetylation levels. (Fig. 2D).

### Validation of acetylated lysine residues based on stable isotope labeling mass spectrometry

To further validate the RCA lysine residues non-enzymatic acetylation by the Ac-CoA, we synthesized a deuterium labeled Ac-CoA (D^2^-Ac-CoA), where three hydrogen atoms of the acetyl group were replaced by the deuterium (D). Based on LC-MS/MS analysis, 6 acetylated RCA lysine residues were characterized from the acetylation assays *in vitro* by the D^2^-Ac-CoA. These are K17, K34, K72, K116, K126 and K164 (Fig. 3A-B; Supplementary Table S3), which are located at the N-terminal sub-domain (Supplementary Fig. S6). Strikingly, the number of the identified acetylated lysine residues *in vitro* was more than expected for the native RCA, which might be owing to the high Ac-CoA concentration or the different biomolecular structure of the RCA between *in vivo* and *in vitro*. This supports the evidence as observed in previous studies that the RCA polymerization could be altered when ATP and RuBP are present in the assay(Mueller-Cajar *et al.*, 2011). This suggests the presence of some other unknown inhibitors in addition to RuBP that could potentially be also involved in the reversible RCA acetylation in the chloroplast.

**Fig. 3.**
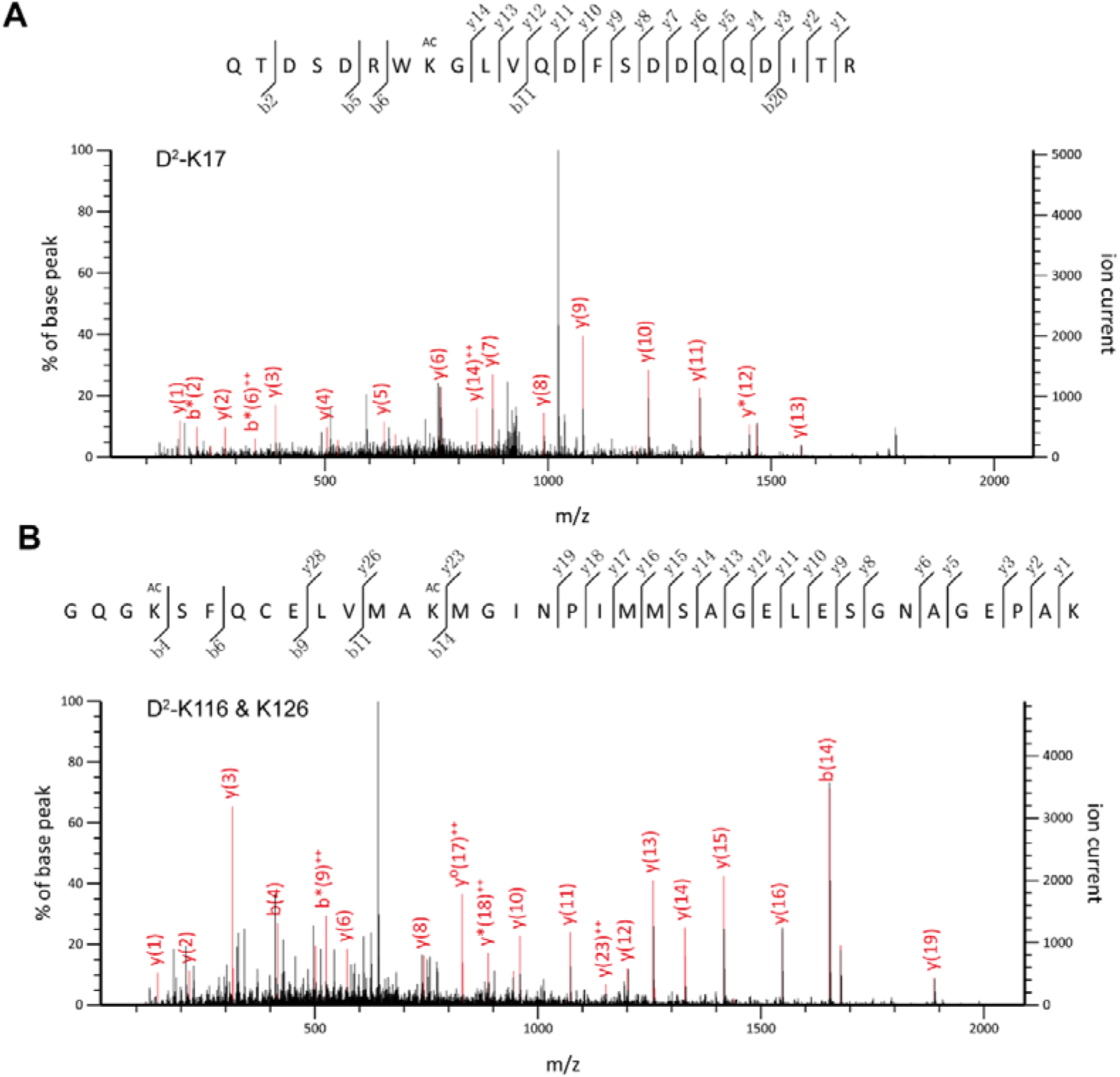
Evaluation of RCA non-enzymatic acetylated by Ac-CoA using deuterium-labeled Ac-CoA (D^2^-Ac-CoA). (**A**-**B**) Mass spectrum of the acetylated lysine residues in tobacco RCA. The latter (RCA) was initially incubated with 0.5 mM D^2^-Ac-CoA for 30 min at 25 °C, then digested and loaded to the LC-MS/MS for further fragmentation (decomposition), analysis and detection. (**A)** Modification of the K17 peptide. (**B)** Modification of the K116 and K126 peptides. The sequences fragments are displayed on the top of each panel.

### The deacetylase inhibitor promotes the RCA acetylation level and decreases its stability

To gain more insights in this subject, we used three known histone deacetylase inhibitors (HDACi), i.e., Nicotinamide (NAM), Trichostatin A (TSA) and sodium butyrate (NaB). Notably, the NAM selectively inhibits the Sirtuin (SIRT) HDACi family, the TSA is a multiple HDACi family inhibitor and the sodium butyrate (NaB) is a histone deacetylase inhibitor. As predicted, the western blots analysis shows that the RCA acetylation level increased in the treated chloroplast with the deacetylase inhibitors if compared with that of the control sample (Fig. 4A), while, interestingly, the RCA content decreased significantly (*P* < 0.05, Fig. 4B). In this regards, it has been earlier revealed that the acetylation features could affect the proteins stability and turnover (Mateo *et al.*, 2009). We thus speculate that the acetylation process might be somehow related to the RCA stability.

**Fig. 4.**
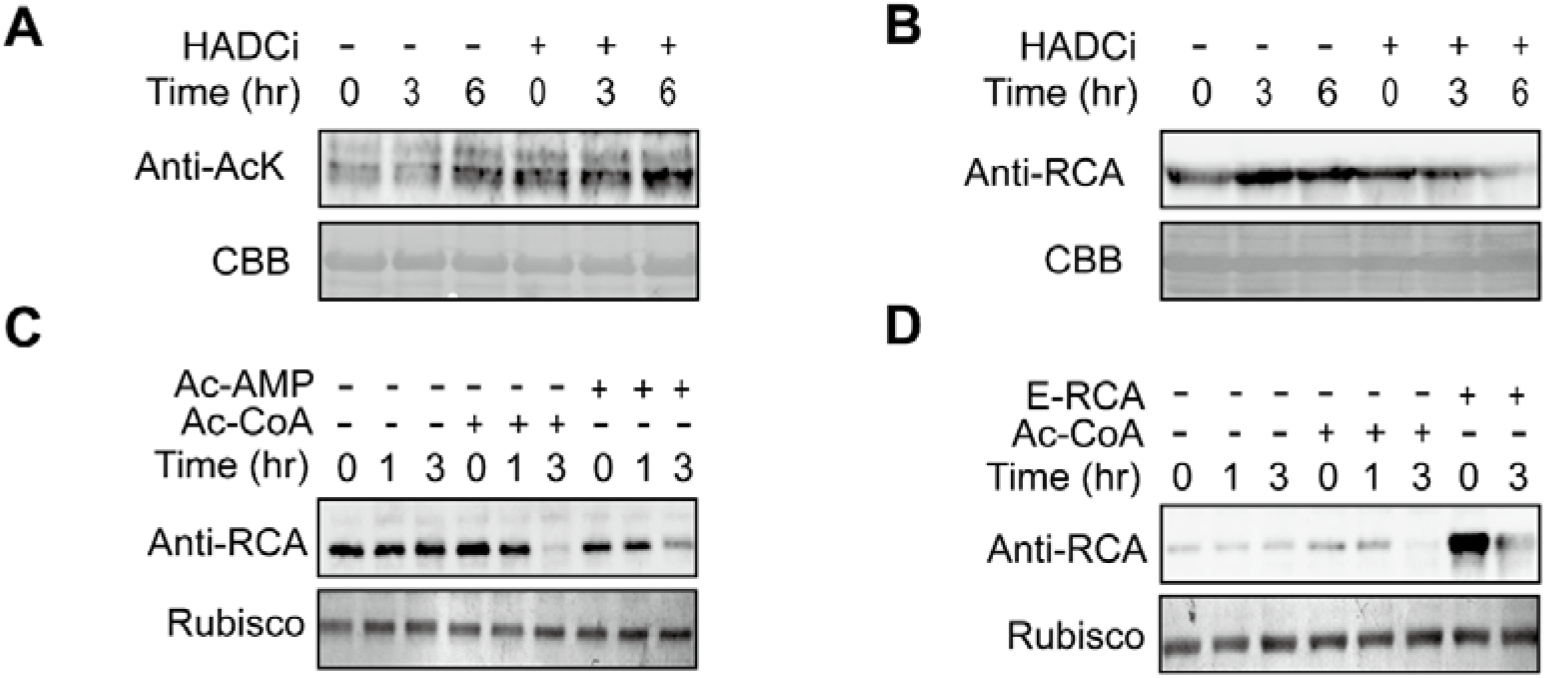
The lysine acetylation adversely affects the RCA stability and activity. (**A**) The lysine acetylation proteins of RCA induced by HDACi for different hours. (**B**) The RCA protein expression levels due to HDACi induced acetylation effects. (**C**) The RCA protein expression levels due to either Ac-AMP or Ac-CoA induced acetylation effects. (**D**) The RCA protein expression levels in E-RCA (expressed in *E. coli*) due to Ac-CoA induced acetylation effects. Intact tobacco chloroplast was treated with HDACi inhibitor, 50 nM TSA, 2 mM NAM and 2 mM NAB (see material and methods for full inhibitors names), then the stroma total soluble proteins were collected at different time-points including 0, 3 and 6 h. The intact tobacco chloroplast was treated with Ac-AMP (10 mM) and Ac-CoA (0.5 mM; **C**) and E-RCA (expressed by *E. coli*) (**D**). For all samples containing HADCi inhibitors, the stroma proteins were collected at 0, 1 and 3 h. The RCA protein levels were determined by western-blot analysis. Data were reproducible and representative of at least 3 independent experiments.

Therefore, we further investigated the changes in RCA content in the chloroplast treated with additional Ac-CoA and Ac-AMP. Consistently, the RCA content also obviously decreased after incubation for 3 hour with AMP or Ac-CoA (Fig. 4C). To prevent the acetylation impact of other proteins which might also directly or indirectly affect the RCA stability in the chloroplast, we added the purified prokaryotic recombinant RCA from *E. coli* (E-RCA), which exhibited much more acetylated lysine residues and RCA acetylation level in the crude chloroplast extracts (Fig. 4D; Supplementary Table S2; Supplementary Table S4). The western-blot analysis demonstrates that the degradation of the recombinant RCA was faster than that of the native one (RCA) in chloroplast, which indicates that the faster RCA degradation is mainly attributed to the extent of RCA acetylation level. These results unravel that the RCA acetylation might serve a sign for the RCA degradation *in vivo*.

Given that the RuBP could inhibit the RCA acetylation by the Ac-CoA, the low RuBP level would result in a higher RCA acetylation level, and in turn leads to a faster RCA degradation in the chloroplast (Figs. 2D and 4C). This statement was also supported by the decreased RCA amount as well as by its (RCA) increased acetylation level in tobacco leaves under a moderate heat stress (Fig. 1B). Actually, it becomes well known that the RCA is a sensitive protein to high temperature (Salvucci *et al.*, 2001). In line with this, our results indicate that the RCA degradation induced following its acetylation may somehow contribute to its thermosensitivity *in vivo* to high temperature.

### RCA degradation following its acetylation is not related to the ubiquitination

In this regards, several reports suggested that the acetylation operation could regulate the protein stability through its ubiquitination *in vivo* (Grice and Nathan, 2016). However, the ubiquitination process is not necessary involved in the protein degradation in the chloroplast (Smalle and Vierstra, 2004; Moon *et al.*, 2004). Consistently, our results showed also that the RCA protein level in the chloroplast was not affected by the ubiquitination upon treatment with 25 μM MG132, an ubiquitin-proteasome inhibitor (Supplementary Fig. S7A-B). In the same trend, it has been reported in a very recent study that the proteins ubiquitination may occur when proteins are exported out of the chloroplast compartment using SP2 and CDC48, a chloroplast-associated protein degradation, CHLORAD (Ling *et al.*, 2019); however, no significant differences were observed in the RCA amount between the mutant *sp2* and Col in Arabidopsis at night (Supplementary Fig. S8). This suggest that some other degradation systems and special protease could regulates RCA degradation in chloroplast as documented previously (Striebel *et al.*, 2009; Fukayama *et al.*, 2012).

Taken together, as shown in Fig. 5, the RCA can be acetylated *in vivo* in the presence of Ac-CoA especially under high temperature. However, RCA acetylation level could be inhibited under light as well as by RuBP. The RCA acetylation has led to a considerable decrease in the ATPase activity and its protein amount. Therefore, the degraded protein, as reflected by the substantial decrease in the RCA protein amount, resulted eventually in a low Rubisco activation activity. Hereby, we deduce that the ubiquitination does not constitute the sole reason standing behind the protein degradation process. Nevertheless, to well understand the mechanism of the RCA degradation further investigations are needed to better decipher the detailed and inter-related biochemical and molecular pathways orchestrating in a well concerted manner this sequential phenomenon (acetylation, ubiquitination and degradation).

**Fig. 5.**
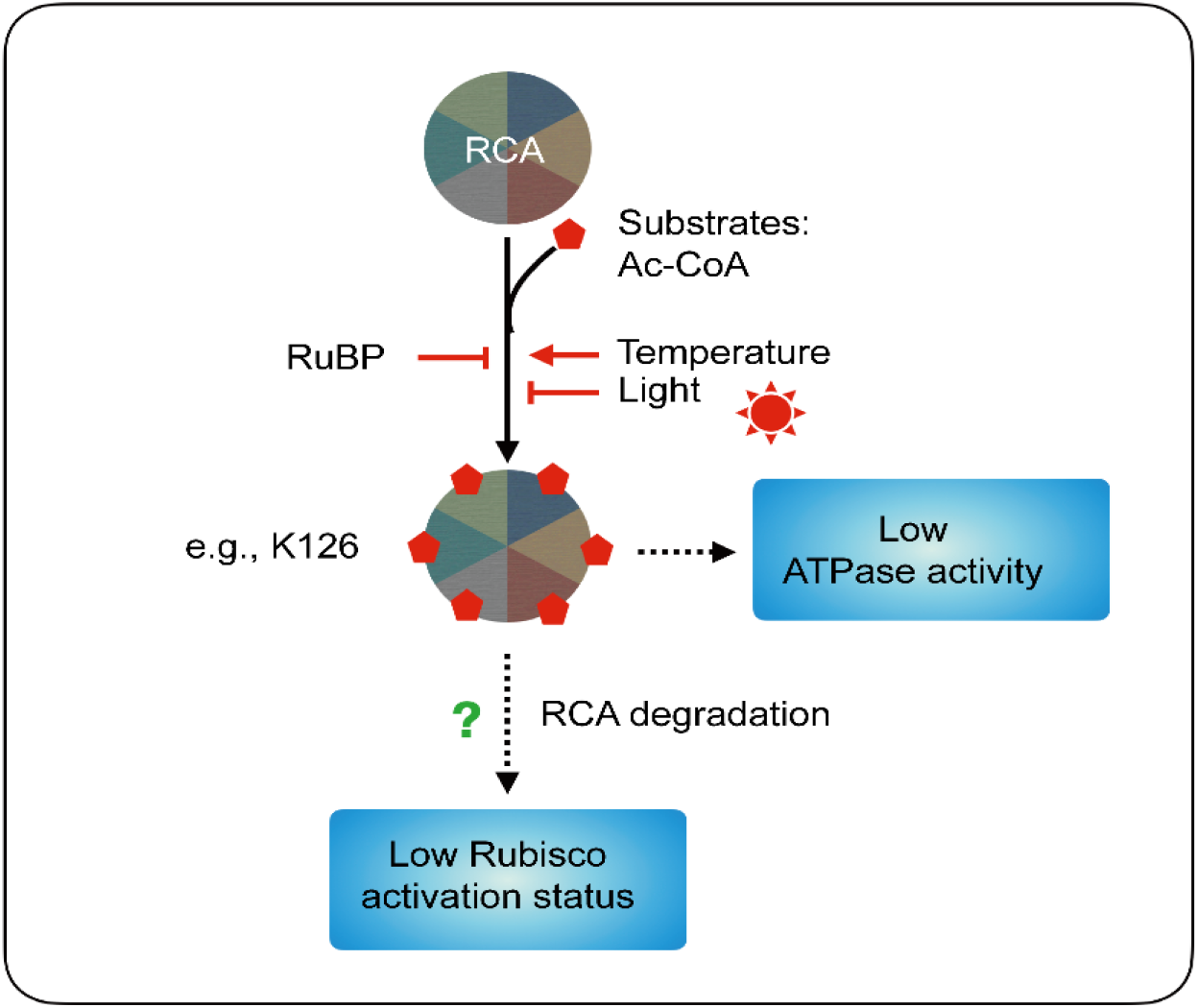
A simplified schematic representative working model depicting the biological relevance and the regulatory mechanisms of the RCA acetylation. The RCA can be acetylated in presence of Ac-CoA *in vivo* and its (RCA) acetylation level could be inhibited by light as well as by RuBP. Thus, the acetylated RCA can slow down its ATPase activity and decreases its protein amount. Therefore, the degraded protein, as reflected by the decreased RCA protein level, may eventually lead to a lower Rubisco activity. Ultimately, we could deduce that the ubiquitination process does not rperent the direct reason standing behind the RCA protein degradation rather it is very likely its acetylation.

## Conclusion

In this study, we revealed that the RCA acetylation level is feasible to temperature and light stimulus, and it could be acetylated non-enzymatically by various metabolites including Ac-CoA through their direct interactions. The acetylated RCA affects its stability, and together with decrease its ATPase activity and Rubisco activation activity. Overall, the finding provides novel insights about the PTMs regulating the RCA function and a model by which the acetylation might react in an enzyme-independent manner in the plant chloroplast.

## Acknowledgments

We thank Professor Qihua Ling for providing *sp2* mutant seeds and professors Wenli Hu, Xiaoyan Xu, and Shimin Zhao for their kindness and patient guidance on the academic knowledge. We also thank all the colleagues Xinyu Liu, Zhaoxue Ma, Xunlin Han, Min Xu, Juan Chen, Zhiyong Xu and Zongzhu Li for generously supporting this study. This work was supported by grants from National Natural Science foundation of China, NSFC (31702019 to H. H., 81803139 to L.Y., 31700201 to M.Q.).

## Author contributions

G.C., X.C., H.M., M.Q. planed and designed the research; M.Q., H.H., X.G., X.F., L.Y., Z.S., A.G., M.W. performed experiments; L.Y., M.Q., J.E., G.C. wrote the manuscript; G.C., M.Q., X.C. supervised the research; L.Y., H.H., contributed equally.

## Data and materials availability

All data is available in the manuscript or the supporting information.

## Reference

Carmo-Silva AE, Salvucci ME. 2013. The regulatory properties of rubisco activase differ among species and affect photosynthetic induction during light transitions. Plant Physiology 161, 1645 LP – 1655.

Chen C, Li C, Wang Y, et al. 2017. Cytosolic acetyl-CoA promotes histone acetylation predominantly at H3K27 in Arabidopsis. Nature Plants 3, 814–824.

Chen J, Wang P, Mi H -l., Chen G-Y, Xu D-Q. 2010. Reversible association of ribulose-1, 5-bisphosphate carboxylase/oxygenase activase with the thylakoid membrane depends upon the ATP level and pH in rice without heat stress. Journal of Experimental Botany 61, 2939–2950.

Cox J, Neuhauser N, Michalski A, Scheltema RA, Olsen J V, Mann M. 2011. Andromeda: A peptide search engine integrated into the MaxQuant environment. Journal of Proteome Research 10, 1794–1805.

Esau BD, Snyder GW, Portis ARJ. 1996. Differential effects of N- and C-terminal deletions on the two activities of rubisco activase. Archives of biochemistry and biophysics 326, 100–105.

Feller U, Crafts-Brandner SJ, Salvucci ME. 1998. Moderately high temperatures inhibit ribulose-1,5-bisphosphate carboxylase/oxygenase (Rubisco) activase-mediated activation of rubisco. Plant Physiology 116, 539 LP – 546.

Finkemeier I, Laxa M, Miguet L, Howden AJM, Sweetlove LJ. 2011. Proteins of diverse function and subcellular location are lysine acetylated in Arabidopsis. Plant Physiology 155, 1779 LP – 1790.

Flecken M, Wang H, Popilka L, et al. 2020. Article dual functions of a rubisco activase in metabolic repair and recruitment to carboxysomes ll article dual functions of a rubisco activase in metabolic repair and recruitment to carboxysomes. Cell 183, 457–473.e20.

Fukayama H, Ueguchi C, Nishikawa K, Katoh N, Ishikawa C, Masumoto C, Hatanaka T, Misoo S. 2012. Overexpression of rubisco activase decreases the photosynthetic co2 assimilation rate by reducing rubisco content in rice leaves. Plant and Cell Physiology 53, 976–986.

Gao X, Hong H, Li WC, Yang L, Huang J, Xiao YL, Chen XY, Chen GY. 2016. Downregulation of rubisco activity by non-enzymatic acetylation of RbcL. Molecular Plant 9, 1018–1027.

Grabsztunowicz M, Koskela MM, Mulo P. 2017. Post-translational modifications in regulation of chloroplast function: recent advances. Frontiers in plant science 8, 240.

Graciet E, Lebreton S, Gontero B. 2004. Emergence of new regulatory mechanisms in the Benson-Calvin pathway via protein-protein interactions: A glyceraldehyde-3-phosphate dehydrogenase/CP12/ phosphoribulokinase complex. Journal of Experimental Botany 55, 1245–1254.

Grice GL, Nathan JA. 2016. The recognition of ubiquitinated proteins by the proteasome. Cellular and molecular life sciences◻: CMLS 73, 3497–3506.

Hasse D, Larsson AM, Andersson I. 2015. Structure of Arabidopsis thaliana Rubisco activase. Acta crystallographica. Section D, Biological crystallography 71, 800–808.

Jensen RG. 2000. Activation of Rubisco regulates photosynthesis at high temperature and CO_2_. Proceedings of the National Academy of Sciences 97, 12937 LP – 12938.

Jiang J, Gai Z, Wang Y, Fan K, Sun L, Wang H, Ding Z. 2018. Comprehensive proteome analyses of lysine acetylation in tea leaves by sensing nitrogen nutrition. BMC Genomics 19, 840.

Kim SY, Harvey CM, Giese J, Lassowskat I, Singh V, Cavanagh AP, Spalding MH, Finkemeier I, Ort DR, Huber SC. 2019. In vivo evidence for a regulatory role of phosphorylation of Arabidopsis Rubisco activase at the Thr78 site. Proceedings of the National Academy of Sciences 116, 18723 LP – 18731.

Kubis SE, Lilley KS, Jarvis P. 2008. Isolation and preparation of chloroplasts from Arabidopsis thaliana plants. Methods in molecular biology (Clifton, N.J.) 425, 171–186.

Kurek I, Chang TK, Bertain SM, Madrigal A, Liu L, Lassner MW, Zhu G. 2007. Enhanced thermostability of Arabidopsis Rubisco activase improves photosynthesis and growth rates under moderate heat stress. The Plant cell 19, 3230–3241.

Kuriata AM, Chakraborty M, Henderson JN, Hazra S, Serban AJ, Pham TVT, Levitus M, Wachter RM. 2014. ATP and magnesium promote cotton short-form ribulose-1,5-bisphosphate carboxylase/oxygenase (Rubisco) activase hexamer formation at low micromolar concentrations. Biochemistry 53, 7232–7246.

Li J, Yokosho K, Liu S, Cao HR, Yamaji N, Zhu XG, Liao H, Ma JF, Chen ZC. 2020. Diel magnesium fluctuations in chloroplasts contribute to photosynthesis in rice. Nature Plants.

Lima BP, Antelmann H, Gronau K, Chi BK, Becher D, Brinsmade SR, Wolfe AJ. 2011. Involvement of protein acetylation in glucose-induced transcription of a stress-responsive promoter. Molecular microbiology 81, 1190–1204.

Ling Q, Broad W, Trösch R, Töpel M, Demiral Sert T, Lymperopoulos P, Baldwin A, Jarvis RP. 2019. Ubiquitin-dependent chloroplast-associated protein degradation in plants. Science 363, eaav4467.

Mateo F, Vidal-Laliena M, Canela N, Busino L, Martinez-Balbas MA, Pagano M, Agell N, Bachs O. 2009. Degradation of cyclin A is regulated by acetylation. Oncogene 28, 2654–2666.

Mischerikow N, Heck AJR. 2011. Targeted large-scale analysis of protein acetylation. Proteomics 11, 571–589.

Moon J, Parry G, Estelle M. 2004. The Ubiquitin-proteasome pathway and plant development. The Plant Cell 16, 3181 LP – 3195.

Mueller-Cajar O, Stotz M, Wendler P, Hartl FU, Bracher A, Hayer-Hartl M. 2011. Structure and function of the AAA ^+^ protein CbbX, a red-type Rubisco activase. Nature 479, 194–199.

O’Leary BM, Scafaro AP, Fenske R, Duncan O, Ströher E, Petereit J, Millar AH. 2020. Rubisco lysine acetylation occurs at very low stoichiometry in mature Arabidopsis leaves: implications for regulation of enzyme function. The Biochemical journal 477, 3885–3896.

Osorio S, Ruan Y-L, Fernie AR. 2014. An update on source-to-sink carbon partitioning in tomato. Frontiers in plant science 5, 516.

Peltier J-B, Cai Y, Sun Q, Zabrouskov V, Giacomelli L, Rudella A, Ytterberg AJ, Rutschow H, van Wijk KJ. 2006. The oligomeric stromal proteome of *Arabidopsis thaliana* chloroplasts. Molecular & Cellular Proteomics 5, 114 LP – 133.

Portis AR. 2003. Rubisco activase – Rubisco’s catalytic chaperone. Photosynthesis Research 75, 11–27.

Post-Beittenmiller D, Roughan G, Ohlrogge JB. 1992. Regulation of plant fatty acid biosynthesis◻: analysis of acyl-coenzyme a and acyl-acyl carrier protein substrate pools in spinach and pea chloroplasts. Plant physiology 100, 923–930.

Ramponi G, Manao G, Camici G. 1975. Nonenzymic acetylation of histones with acetyl phosphate and acetyl adenylate. Biochemistry 14, 2681–2685.

Robinson SP, Portis AR. 1989. Adenosine triphosphate hydrolysis by purified rubisco activase. Archives of Biochemistry and Biophysics 268, 93–99.

Salvucci ME, Crafts-Brandner SJ. 2004. Relationship between the heat tolerance of photosynthesis and the thermal stability of Rubisco activase in plants from contrasting thermal environments. Plant Physiology 134, 1460 LP – 1470.

Salvucci ME, Osteryoung KW, Crafts-Brandner SJ, Vierling E. 2001. Exceptional sensitivity of Rubisco activase to thermal denaturation in vitro and in vivo. Plant physiology 127, 1053–1064.

Shen JB, Orozco EMJ, Ogren WL. 1991. Expression of the two isoforms of spinach ribulose 1,5-bisphosphate carboxylase activase and essentiality of the conserved lysine in the consensus nucleotide-binding domain. The Journal of biological chemistry 266, 8963–8968.

Smalle J, Vierstra RD. 2004. The ubiquitin 26S proteasome proteolytic pathway. Annual Review of Plant Biology 55, 555–590.

Spreitzer RJ, Salvucci ME. 2002. Rubisco: structure, regulatory interactions, and possibilities for a better enzyme. Annual review of plant biology 53, 449–475.

Stotz M, Mueller-Cajar O, Ciniawsky S, Wendler P, Hartl FU, Bracher A, Hayer-Hartl M. 2011. Structure of green-type Rubisco activase from tobacco. Nature Structural & Molecular Biology 18, 1366–1370.

Striebel F, Kress W, Weber-Ban E. 2009. Controlled destruction: AAA^+^ ATPases in protein degradation from bacteria to eukaryotes. Current opinion in structural biology 19, 209–217.

Tumaney AW, Ohlrogge JB, Pollard M. 2004. Acetyl coenzyme A concentrations in plant tissues. Journal of plant physiology 161, 485–488.

Wagner GR, Payne RM. 2013. Widespread and enzyme-independent Nε-acetylation and Nε-succinylation of proteins in the chemical conditions of the mitochondrial matrix. The Journal of biological chemistry 288, 29036–29045.

Wang Q, Zhang Y, Yang C, et al. 2010. Acetylation of metabolic enzymes coordinates carbon source utilization and metabolic flux. Science 327, 1004 LP – 1007.

Weinert BT, Iesmantavicius V, Wagner SA, Schölz C, Gummesson B, Beli P, Nyström T, Choudhary C. 2013. Acetyl-phosphate is a critical determinant of lysine acetylation in E. coli. Molecular cell 51, 265–272.

Wu X, Oh M-H, Schwarz EM, Larue CT, Sivaguru M, Imai BS, Yau PM, Ort DR, Huber SC. 2011. Lysine acetylation is a widespread protein modification for diverse proteins in Arabidopsis. Plant Physiology 155, 1769 LP – 1778.

Yamori W, Masumoto C, Fukayama H, Makino A. 2012. Rubisco activase is a key regulator of non-steady-state photosynthesis at any leaf temperature and, to a lesser extent, of steady-state photosynthesis at high temperature. The Plant journal[: for cell and molecular biology 71, 871–880.

Zhang N, Portis ARJ. 1999. Mechanism of light regulation of Rubisco: a specific role for the larger Rubisco activase isoform involving reductive activation by thioredoxin-f. Proceedings of the National Academy of Sciences of the United States of America 96, 9438–9443.

Zhou L-J, Li Y-Y, Zhang R-F, Zhang C-L, Xie X-B, Zhao C, Hao Y-J. 2017. The small ubiquitin-like modifier E3 ligase MdSIZ1 promotes anthocyanin accumulation by sumoylating MdMYB1 under low-temperature conditions in apple. Plant, cell & environment 40, 2068–2080.

